# Analysis of the impact of molecular motions on the efficiency of XL-MS and the distance restraints in hybrid structural biology

**DOI:** 10.1101/379289

**Authors:** Vladimir Svetlov, Evgeny Nudler

## Abstract

Covalent cross-link mapping by mass spectrometry (XL-MS) is rapidly becoming the most widely used method of hybrid structural biology. We investigated the impact of incremental variations of cross-linker length have on the depth of XL-MS interrogation of protein-protein complexes, and assessed the role molecular motions in solution play in generation of cross-link-derived distance restraints. Supplementation of a popular NHS-ester cross-linker, DSS, with 2 reagents shorter or longer by CH2-CH2, increased the number of non-reductant cross-links by ^~^50%. Molecular dynamics simulations of these cross-linkers revealed 3 individual, partially overlapping ranges of motion, consistent with partially overlapping sets of cross-links formed by each reagent. Similar simulations elucidated protein fold-specific ranges of motions for the reactive and backbone atoms from rigid and flexible target domains. Together these findings create a quantitative framework for generation of cross-linker- and protein fold-specific distance restraints for XL-MS-guided protein-protein docking.

---

Covalent cross-link mapping assisted by mass spectrometry (XL-MS, also CXMS, and CLMS) is a low-resolution method of structural interrogation of proteins and protein-protein complexes (PPCs), which employs chemical probing of reactive groups on protein surfaces with bi-functional covalent cross(X)-linkers^1-5^. Other methods focusing on detection of surface/solvent-accessible amino acids, such as hydrogen-deuterium exchange or photo-oxidation, return information about single residues located either to the surface or the interior of the PPC^6,7^. XL-MS by the virtue of mass spectrometry-assisted identification of covalently X-linked pairs of amino acids provides a set of pairwise distance restraints, dependent on the length of the X-linker, the structure and the conformational dynamics of a given PPC. These distance restraints in turn can assist with interpretation of crystallographic and cryo-EM experimental densities, as well as guiding protein folding and docking computations^8-12^.

Lagging behind X-ray crystallography and cryo-EM in resolution and completeness of structural information, XL-MS has a set of distinct advantages over them: it is less sensitive to the purity and homogeneity of the PPC sample, and utilizes broadly distributed instruments, whose high sensitivity accounts for lower sample consumption. Even more importantly, by utilizing cell-permeable X-linkers XL-MS can simultaneously discover the composition and report on the structure of PPCs *in vivo*, using linear and X-linked peptides, respectively^13^. For a long time since publication of the first experimental application of XL-MS^14^ this technique occupied a relatively small niche in structural biology, predominantly populated by proof-of-principle publications^1^. The past obstacles to the broad utilization of XL-MS can be traced to the highly labor-intensive and idiosyncratic manual validation of the spectral data, the focus on custom-made functionalized reagents, and the relative scarcity and low sensitivity of mass spectrometers^1,15,16^.

Recently XL-MS use has been steadily expanding, propelled by the development of automatic (score-driven) discovery algorithms^17-19^, and the proliferation of the highly sensitive Orbitrap instruments, greatly facilitating data-dependent data acquisition and thus discrimination between linear and X-linked peptides based on their charge^20^. The portability of the XL-MS pipeline has been further improved by shifting the emphasis to the commercially available amino-reactive NHS-esters, such as disuccinimidyl suberate (DSS) and bis(sulfosuccinimidyl)suberate (BS3)^21^. These reagents offer numerous advantages in addition to their broad availability: their chemical properties and reactivity are studied in far greater detail than the rest of X-linker reagents; at near neutral pH (6.5-8.0) NHS-esters react predominantly with Lys residues (and the N-terminal amines), allowing for the formulation of robust discovery rules^21-25^; the relatively high abundance of Lys residues and their preferential localization to protein surfaces (6 and 99%, respectively)^26,27^ provides for the high abundance of X-links; X-linker aliphatic spacer doesn’t excessively fragment during sample preparation or processing (thereby reducing sample loss and improving discovery rate)^1,28-31^.

The biggest remaining obstacle to the realization of XL-MS full potential as a hybrid structural biology methodology lies in the lack of the detailed understanding of the origins and the nature of the distance restraints it produces. The original idea of the covalent X-link serving as a Euclidean “molecular ruler”^28^ connecting two residues has been transformed into the one of a X-linker acting as a upper-limited proximity sensor, yet the exact boundaries of this proximity are still poorly elucidated^32^. In this work we report a large-scale XL-MS interrogation of a multi-subunit, high-value target PPC (*E. coli* RNA polymerase^33^ (RNAP)) with 3 different X-link reagents of incrementally varied length, interpreted in the framework of molecular dynamics (MD) simulations of the relevant distance restraints^34^. The report concludes with recommendations on incorporating XL-MS-derived distance restraints into PPC docking applications and improving the depth of XL-MS experiments by combining X-linkers of different length.

## Results

Two main approaches exist for determining the boundary of distance restraints set by covalent X-links. The first, geometric approach defines the upper limit of such boundary as the Euclidean length of the fully extended X-link; in case of Lys-Lys X-links this is calculated as NZ(=N_ζ_)-NZ, CB(=C_β_)-CB, or CA(=C_α_)-CA distance (**Fig. 1A**). The second approach derives distance restraints from empirical XL-MS data by measuring distances between X-linked residues in experimental (X-ray, and cryo-EM) protein structures^35,36^. In order to inject physical realism into determination of such measurements, Aebersold and colleagues introduced a non-Euclidean Solvent-Assessible Surface Distance (SASD)^37^. Unfortunately, in XL-MS experiments a fraction of X-links is routinely scored as “aberrant” due their apparent length exceeding that of Euclidean upper limit. Moreover, shorter X-linkers tend to produce 2-4 times fewer X-links than the longer ones, although the average length of the X-links in both groups differ only marginally^1^. The uncertainty regarding the “real” length of the experimental X-links and the chances of poisoning the dataset with “aberrant” ones continue to impede successful application of XL-MS in structural biology. X-link-guided protein-protein docking is particularly vulnerable to the uncertainty of the distance restraints. Published reports vary in the estimates of the impact XL-MS-derived restraints had on docking efficiency, and whether SASD performed better than the simple Euclidean distance^9,29,32,35,38,39^.

**Figure 1.**
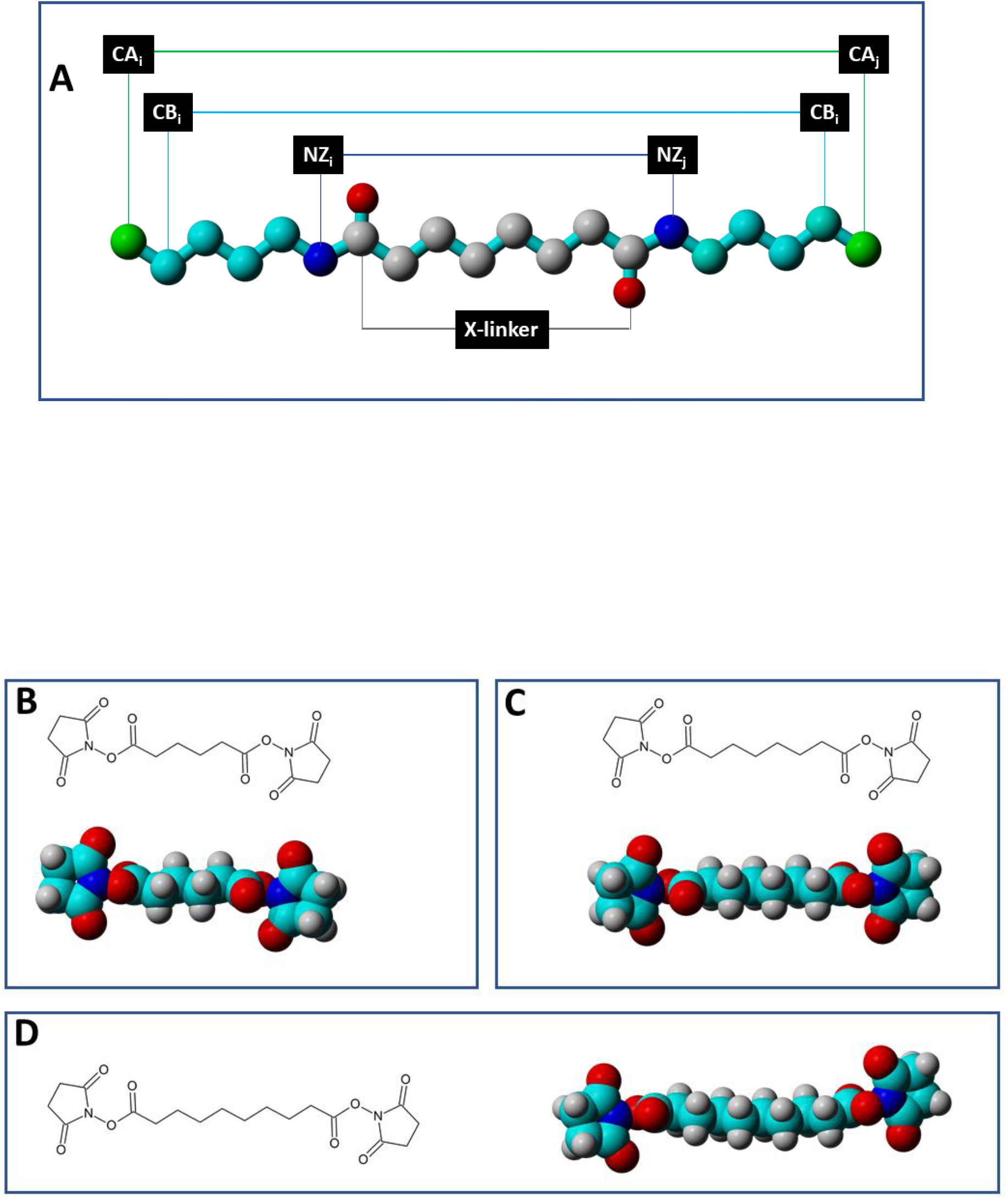
Structures of the NHS-ester X-linkers DSM4, DSM6, and DSM8. **A.** Schematic of DSM6-derived X-link between 2 Lys residues (only CA-NZ chains are shown) defining CA-CA, CB-CB, NZ-NZ distances. **B.** Structural formula and the space-fill model of DSM4. **C.** Structural formula and the space-fill model of DSM6. **D.**

We posit that, once X-link discovery protocol has been validated through structural benchmarking and manual inspection of the spectra, it does not produce any “aberrant” X-links; instead all the high-confidence X-links report on the state of the target PPC in solution. The source of the “aberrant” X-links can be then traced to the aberrant states of the PPC, namely to the presence of misfolded PPC components (deviations from the monomer structure), and the oligomerization of the PPC itself (deviations from the oligomer structure). Although non-specific aggregation of proteins in solution does not produce high-abundance reproducible X-links (albeit it does reduce the overall discovery rate), specific aggregation (dimerization, etc) can not only reduce the prominence of structure-conforming X-links, but also introduce an oligomer-specific X-linked pairs which would be inadvertently scored as “aberrant” in the monomer structure. The aggregation of the sample in XL-MS experiments can be monitored by dynamic light scattering (DLS)^40^, both before and during X-linking, and its negative impact minimized by the optimization of the experimental conditions and/or removal of the aggregated sample from the pipeline.

Derivation of the distance restraints from the length of the X-linker or based on the location of the X-linked residues in the static crystallographic or cryo-EM models fails to account for the conformational flexibility of the X-linker and mobility of the PPC in solution. Thermal molecular motions, largely suppressed in X-ray crystallography and cryo-EM, effectively convert both types of molecules into variously populated ensembles, thereby introducing uncertainty into distance restraints. First evaluation of the conformational flexibility of X-linkers assessed by MD simulations was provided by Houk and colleagues, indicating that DSS in solution populates a number of states, significantly compacted relative to its extended conformation^41^. Unfortunately, this work has not been expanded much further. Conformational mobility of the PPCs has even greater impact on the derivation of the distance restraints. It can be divided into two types of motion: thermal and concerted. The first type comprises motions of the Lys side chains relative to the backbone (NZ_i_-CA_i_ vector), motions of the backbone within given domain (CA_i_-CA_j_), and motions of the domains relative to each other. These can be assessed by subjecting an atomic resolution structure of PPC to MD simulations. The second type of motion comprises non-random, large-scale, and well-defined movements of domains relative to each other as a result of binding to ligands, enzyme’s mechano-chemical cycling, etc^42,43^. Such concerted motions can not be fully explored by the MD simulations, without the input of information about the starting and final conformations, and are outside the scope of this work.

In order to explore molecular motions of covalent X-linkers and the residues they react with we have subjected aliquots of a highly monodisperse, monomodal (8.5% polydispersity, 98% of mass estimated as ~385 kDa (vs theoretical mass of 389 kDa)) preparation of *E. coli* RNAP to X-linking with a panel of reagents, comprised of Adipic acid bis[N-hydroxysuccinimide ester] (DS**M4**, featuring **4 methylene** groups in its spacer), DSS (DSM6), and Disuccinimidyl sebacate (DSM8) (**Fig. 1B-D**). XL-MS data for each X-linker was filtered by false discovery rate (<5%) and e-value (≤0.001)^44^, and pooled from several individual experiments to yield 3 datasets containing ~5 000 individual spectra/precursors (5021, 5130, and 5156, respectively) (**Supplementary Table 1**).

### Combining X-linkers of different length dramatically improve the coverage of XL-MS dataset

RNAP contains 202 Lys residues, 197 (97.5%) of which were discovered to form X-links with one or more X-linkers. All 3 Lys present in the smallest RNAP subunit, RpoZ, formed all 3 types of X-links, whereas the larger subunits exhibited minor differences in reactivity: out of 16 Lys residues present in RpoA 12 were found in peptides X-linked by all 3 reagents, 1 was found in DSM6- and DSM8-derived data, 1 – only among DSM4 X-links, 1 – only among DSM8 X-links, and 1 was not discovered in any of the X-linked peptides. Similarly, 85 out total 87 Lys residues present in the largest subunit, RpoC, were found in the X-linked peptides, 75 formed X-links of all 3 types, 2 were found only in DSM4 dataset, 1 – in DSM4 and DSM8 datasets, 5 – in DSM6 and DSM8 datasets, and 2 – in DSM8 dataset only. Altogether the residues reacting only with subsets of X-linkers are responsible for less than 1% of total X-links and therefore can not explain the 2-3 fold difference in the number of X-links formed by shorter and longer X-linkers^1^.

Analysis of the abundance (the number of precursors per each unique X-link) of X-links distributed among the 3 datasets revealed that the shortest X-linker, DSM4, tends to form a few highly abundant X-links, whereas the majority of X-links have a small (less than 0.1% of the total) number of precursors, drastically reducing the chances of their discovery in each individual experiment (usually ≤1000 precursors). 783 DSM4-derived X-links have a total of 5021 corresponding precursors, with 3 most abundant ones represented by 318, 254, and 131 precursors (14%), and top 20 X-links – by 1477 precursors (29.4%). In comparison, abundance of X-links generated by DSM6 and DSM8 is distributed more evenly: the top 3 X-links in each case account for 115/98/83 and 93/65/65 precursors, respectively, with top 20 being represented by 19.7% (DSM6) and 18.2% (DSM8) of total number of precursors. Since in each individual XL-MS experiments the data file contains 500-1000 X-linked precursors, and at least 1 precursor is required for X-link discovery, the highly uneven sampling by DSM4 is likely to result in a lower number of unique X-links, compared to that of longer X-linkers, DSM6 and DSM8.

Together the three X-linkers lead to discovery of 1315 non-redundant X-links, including 481 inter-subunt ones (**Fig. 2**). DSM4 accounted for 783 total X-links, 167 of them not found in DSM6- and DSM8-derived datasets. Similarly, DSM8 experiments yielded 793 X-links, among which 121 were unique to this reagent. The most commonly used X-linker, DSM6 (DSS) accounted for 907 X-links, 210 of them being DSM6-specific. In terms of the depth of coverage the use of DSM4 and DSM8 improves the XL-MS discovery rate by 44.9%, compared to DSM6 alone. This increase is greater than those reported in the literature, and does not require switching to a different X-linker chemistry or altering the conditions of X-linking and/or sample processing^29^.

**Figure 2.**
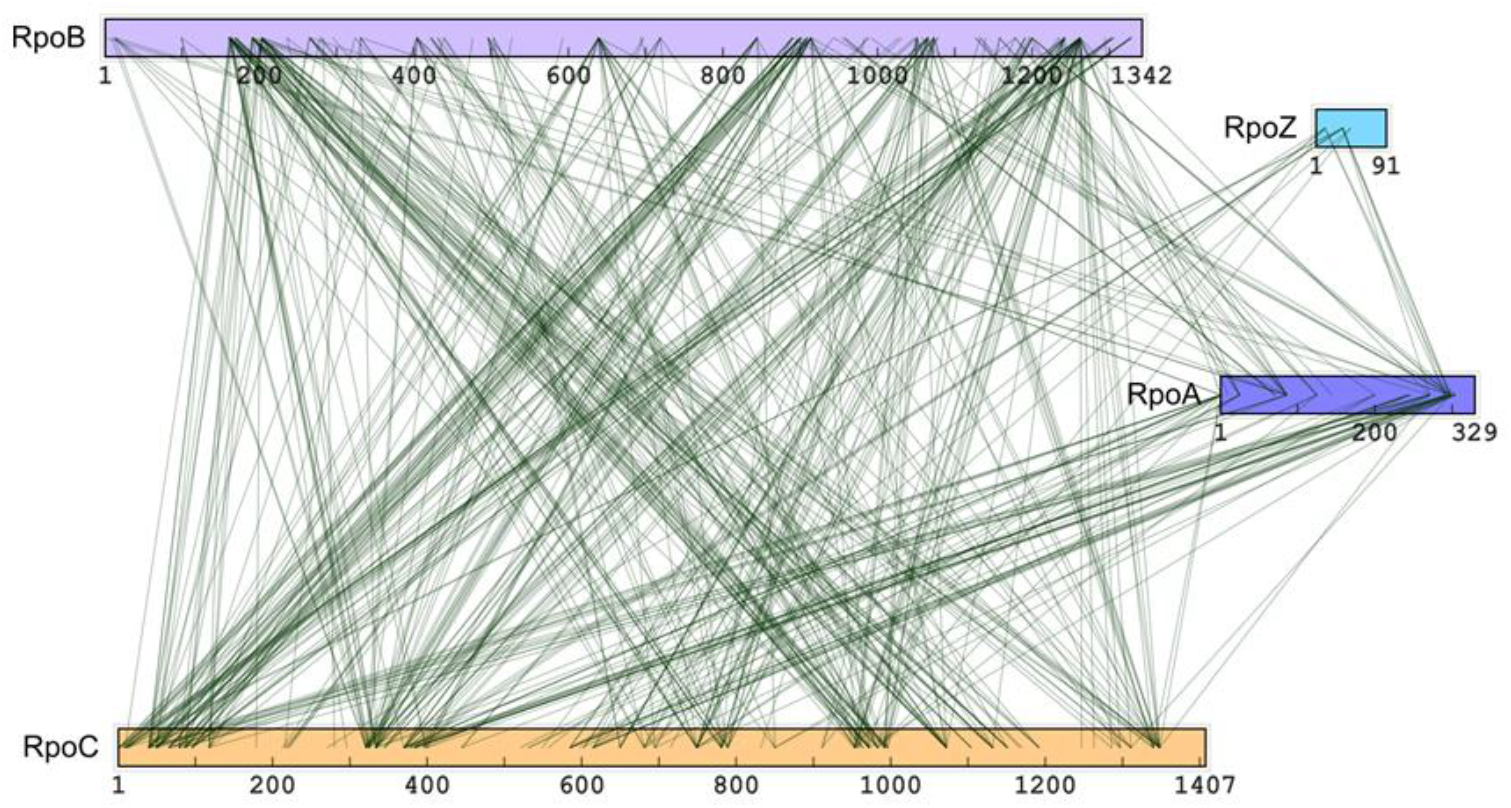
Molecular dynamics simulations of DSM4, DSM6, and DSM8 spacers. Conformational dynamics of NHS-ester X-linkers DSM4/6/8 reflected as temporal fluctuations of the interatomic distances between the first and the last carbon atoms of their respective spacer.

### X-linkers exhibit significant conformational flexibility in solution depending on the length of the spacer

The extended Euclidean length of the X-linker provides the theoretical upper limit of the X-link distance restraint, but since the length of the longer X-link fully encompasses that of the shorter one, it fails to account for the unique X-links formed by the latter. Following the example of Houk and colleagues and our own MD simulation of BS3^45^, we have carried out 25 ns-long full-atom MD simulations of DSM4, DSM6, and DSS8 in explicit water^41^. The spacer length was recorded as [C4;C9] (DSM4), [C4; C11] (DSM6), and [C4,C13] (DSM8) (**Supplementary Tables 2-4**). Consistent with the previous reports and the notion of X-linkers conformational flexibility, these simulations revealed that in solution X-linkers adopt a range of conformations shorter than their Euclidean length (**Fig. 3**). DSM4 linker average length is 5.8 Å vs extended 6.3 Å, ranging from 3.6 to 6.6 Å. DSM6 range of conformations, 4.1-9.1 Å, overlaps with that of DSM4, its average length being 7.7 Å compared to the Euclidean one of 8.7 Å. Similarly, the longest X-linker in the series, DSM8, has an average length of 9.9 Å, down from the Euclidean of 11.2 Å. Its range of 5.5-11.7 Å partially overlaps not only with that of DSM6, but also of DSM4. This partial overlap is consistent with a subset of X-links common to all 3 X-linkers, as well as with presence of unique, X-linker-specific X-links in each dataset. In addition, these simulations provide first principles, quantitative estimates of the distance restraints (lower and upper limits, as well as the median (“target”) distance) imposed by the X-linkers for use in X-link-guided protein-protein docking.

**Figure 3.**
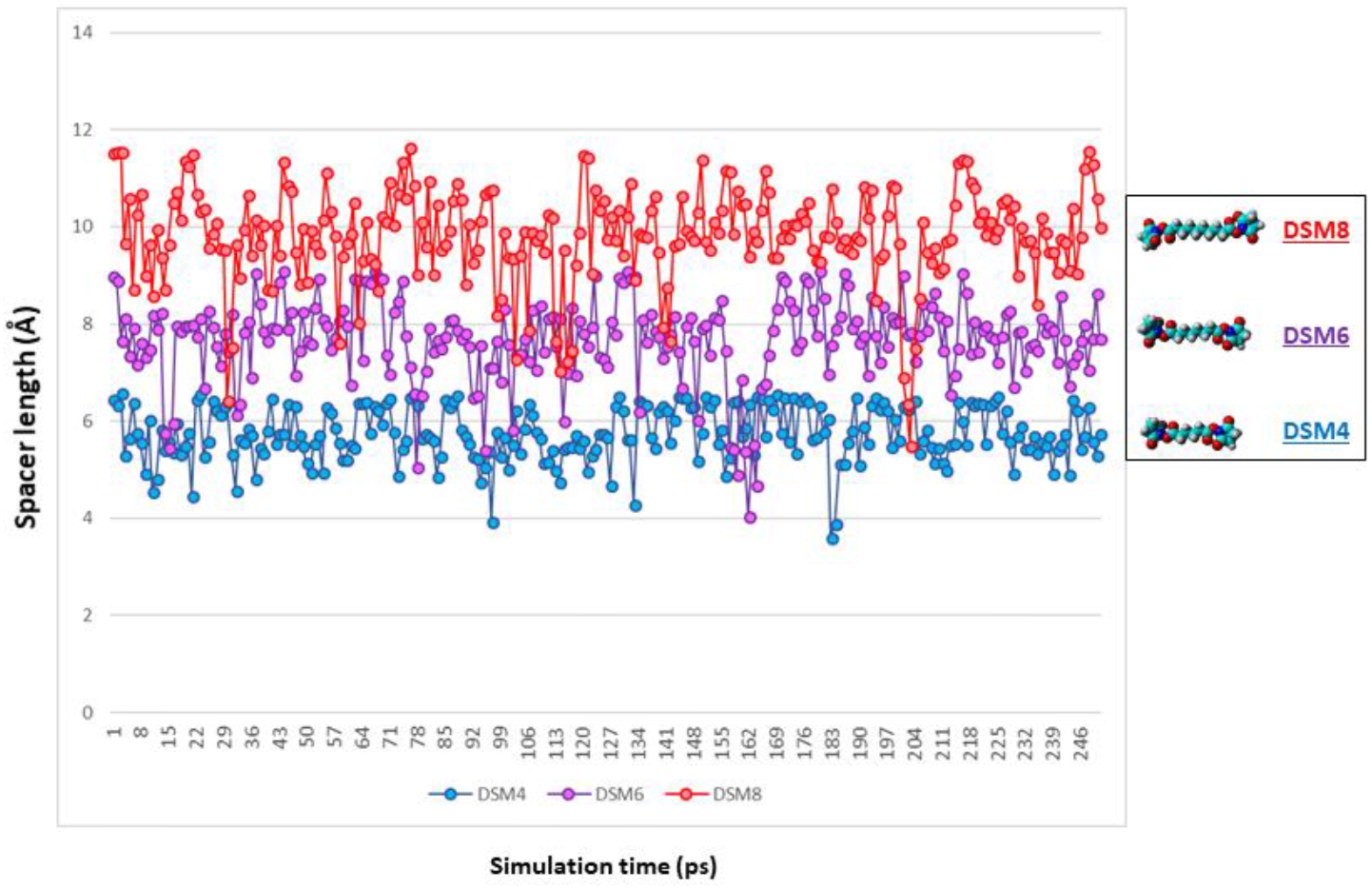
Network view of 481 non-redundant inter-protein X-links between subunits of *E. coli* RNAP subunits. Non-redundant inter-protein X-links aggregated from DSM4/6/8 datasets were mapped to the sequences of RNAP subunits. RpoA subunit present as a dimer in RNAP is shown here as a single copy due to the fact that XL-MS/MS does not permit unambiguous assignment of X-links within homo-oligomeric PPCs.

### The distance between X-linkable amino acids in solution is determined by the fold of the protein and the thermal motions within the protein domains

Conformational dynamics is often invoked in treatment of X-links as distance restraints, but it has not been addressed explicitly and/or quantitatively. Instead, in order to account for this uncertainty in data, the upper limit of X-link restraints has been arbitrarily extended above its Euclidean length. Such indiscriminate, “one-size-fits-all” relaxation of the XL-MS-derived distance restraints is the likely reason why in some cases these restraints failed to significantly improve the quality of the docking models. We have reasoned that the conformational mobility of the X-link-reactive residues is affected by the type of the protein fold they are embedded in, thereby allowing assigning fold-specific restraints values to the individual X-links in the experimental data^46^.

To contrast the fold-dependent motions within protein domains we have extracted the atomic coordinates of 2 domains from the structure of *E. coli* RNAP (4lk1 (**Fig. 4A**)): a rigid α-helical hairpin (RpoB residues 936-1046 (**Fig. 4B**)) and a flexible loop domain (RpoC residues 36-103 (**Fig. 4C**)), and subjected them to the full-atom, explicit water MD simulations as described above. The distances between the actual X-linkable amines (NZ_i_-NZ_j_) and the backbone carbon atoms (CA_i_-CA_j_) were recorded over the course of 20 ns simulations (Supplementary Tables 5 and 6). CA_i_-CB_i_ were also recorded and showed <0.15 Å variation (**Supplementary Table 5**), arguing that behavior of CB_i_-CB_j_ pairs is essentially congruent to that of CA_i_-CA_j_ ones. Consistent with the general expectations residues in the unstructured flexible loop exhibited broader range of motions than those from the α-helical hairpin, with NZ-NZ atoms showing larger displacement relative to the starting structure than CA-CA pairs. Residues forming experimental intra-loop X-links, RpoC50-RpoC66 and RpoC50-RpoC87, started with NZ-NZ distances of 10.0 and 8.9 Å, respectively, but in the course of the MD simulation reached minimum distances of 4.7 and 5.2 Å, and maximum distances of 32.3 and 25.3 Å, with median values of 18.7 and 13.4 Å, respectively (**Fig. 5A**). CA-CA trajectories for the same pairs of X-linkable residues exhibited lower levels of displacement from the starting distances of 10.8 and 14.8 Å, respectively, to the minimum of 9.0 and 10.3 Å and maximum of 27.2 and 22.3 Å (**Fig. 5A**). The median distances were 16.3 and 15.2 Å. The range of motions for residues embedded in the α-helical hairpin found among experimental X-links were as follows (**Fig. 5B**). CA-CA distances for RpoB941-RpoB1035 and RpoB988-RpoB1007 at the start of the simulations were 14.9 and 19.7 Å, respectively. Over 20 ns of simulation time they reached minimum values of 14.1 and 16.3 Å, maximum values of 24.3 and 20.6 Å, with respective median distances of 17.0 and 17.9 Å. NZ-NZ distance variations were larger than those of CA-CA within the same domain, but less dramatic compared to NZ-NZ displacements in the unstructured loop: min-to-max displacements of 9.9 and 13.4 Å (RpoB988-RpoB1007 and RpoB941-RpoB1035) vs 20.1 and 27.6Å (RpoC50-RpoC87 and RpoC50-RpoC66) (**Fig. 6A**). This trend extends to the displacement of CA atoms in the rigid and flexible domains, notably of lower amplitude compared to that of NZ ones (**Fig. 6B**). These data provide quantitative computational framework for the explicit treatment of XL-MS-derived distance restraints in docking simulations and other hybrid methods of structural biology^10,11,35,39,46,47^. They allow to transition from an indiscriminate single distance restraint^28^ to a set of protein/domain-specific ones, and support the notion that the use of CA-CA restraints reduces the uncertainty arising due to molecular thermal motion, as compared to the more intuitive NZ-NZ restraints.

**Figure 4.**
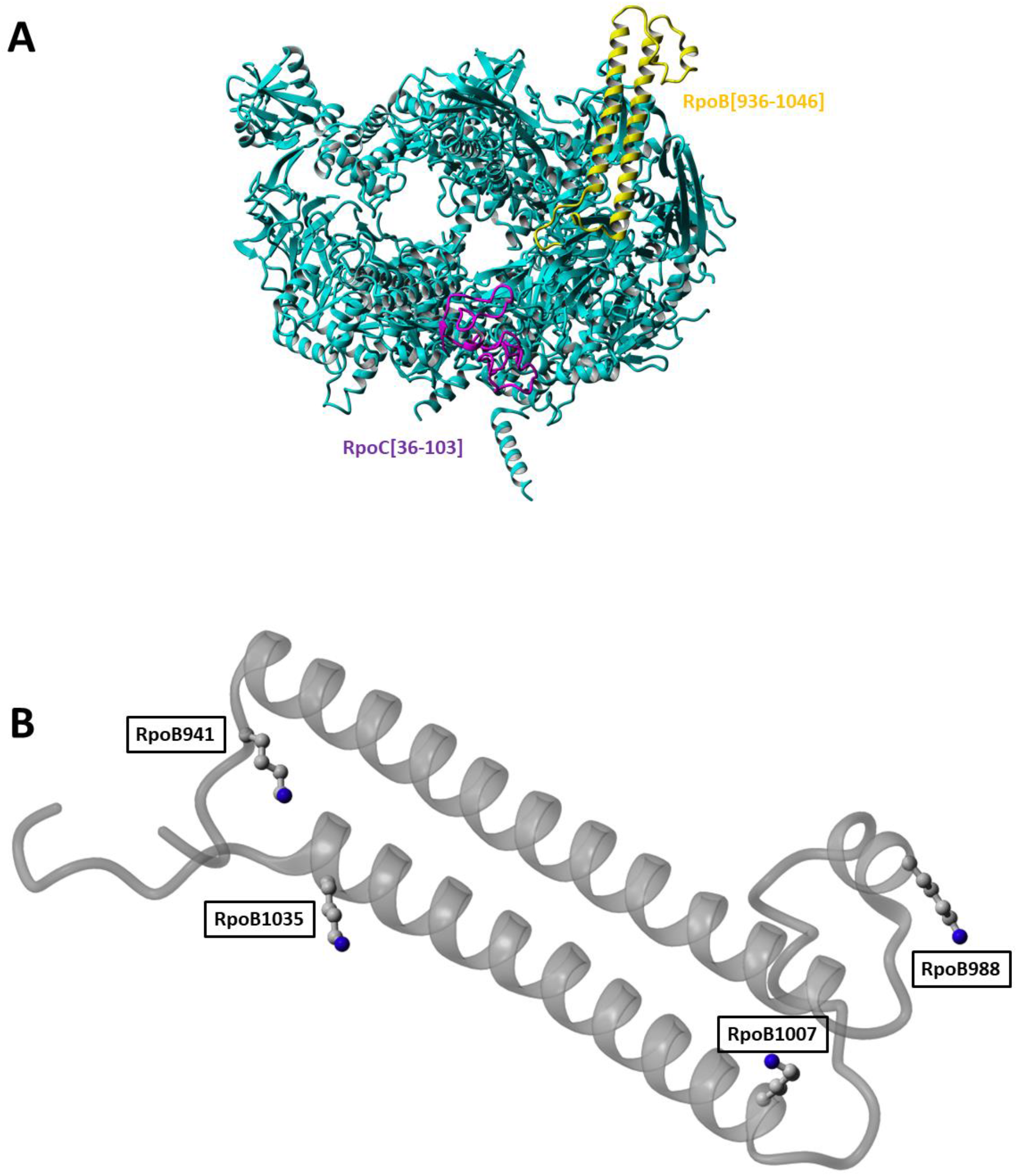
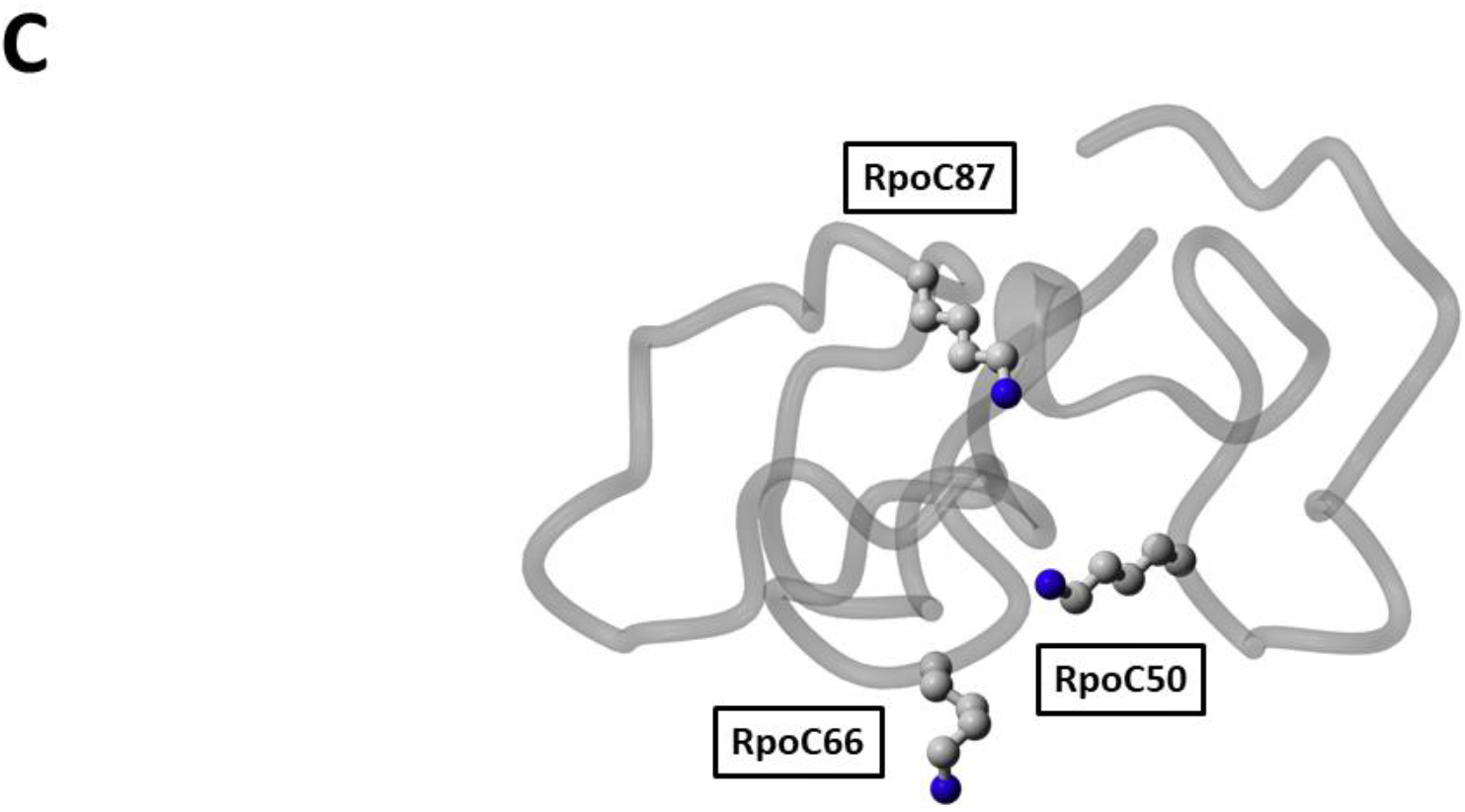
Flexible and rigid domains in the *E. coli* RNA polymerase. **A.** Structure of RNA polymerase core (4lk1), teal, with α-helical hairpin (RpoB(936-1046), yellow), and unstructured loop (RpoC(36-103), purple) domains. **B.** α-helical hairpin (RpoB(936-1046), gray ribbon) with X-linkable Lys residues 941, 988, 1007, and 1035 (ball and stick, gray, NZ atoms highlighted in blue). **C.** unstructured loop (RpoC(36-103), gray ribbon) with X-linkable Lys residues 50, 66, and 87 (ball and stick, gray, NZ atoms highlighted in blue).

**Figure 5.**
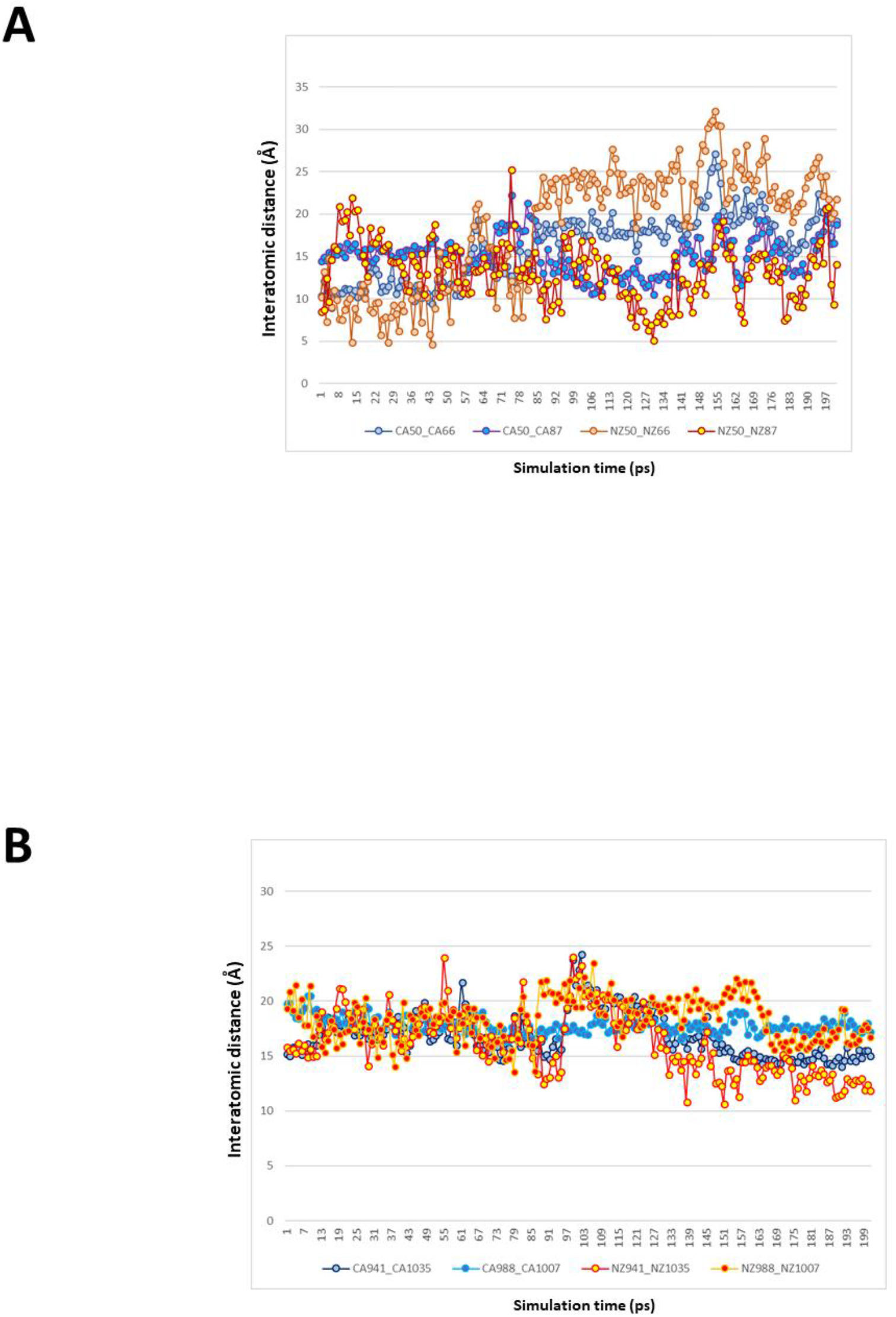
Comparison of the conformational dynamics in rigid and flexible protein domains. **A.** Interatomic distances between backbone CA-CA (CA50-CA66, CA50-CA87) exhibit smaller amplitude than side chain NZ-NZ (NZ50-NZ66, NZ50-NZ66) in the unstructured loop domain (RpoC(36-103)). **B.** Interatomic distances between backbone CA-CA (CA941-CA1035, CA988-CA1007) exhibit smaller amplitude than side chain NZ-NZ (NZ941-NZ1035, NZ988-NZ1007) in the rigid α-helical hairpin (RpoB(936-1046) domain.

**Figure 6.**
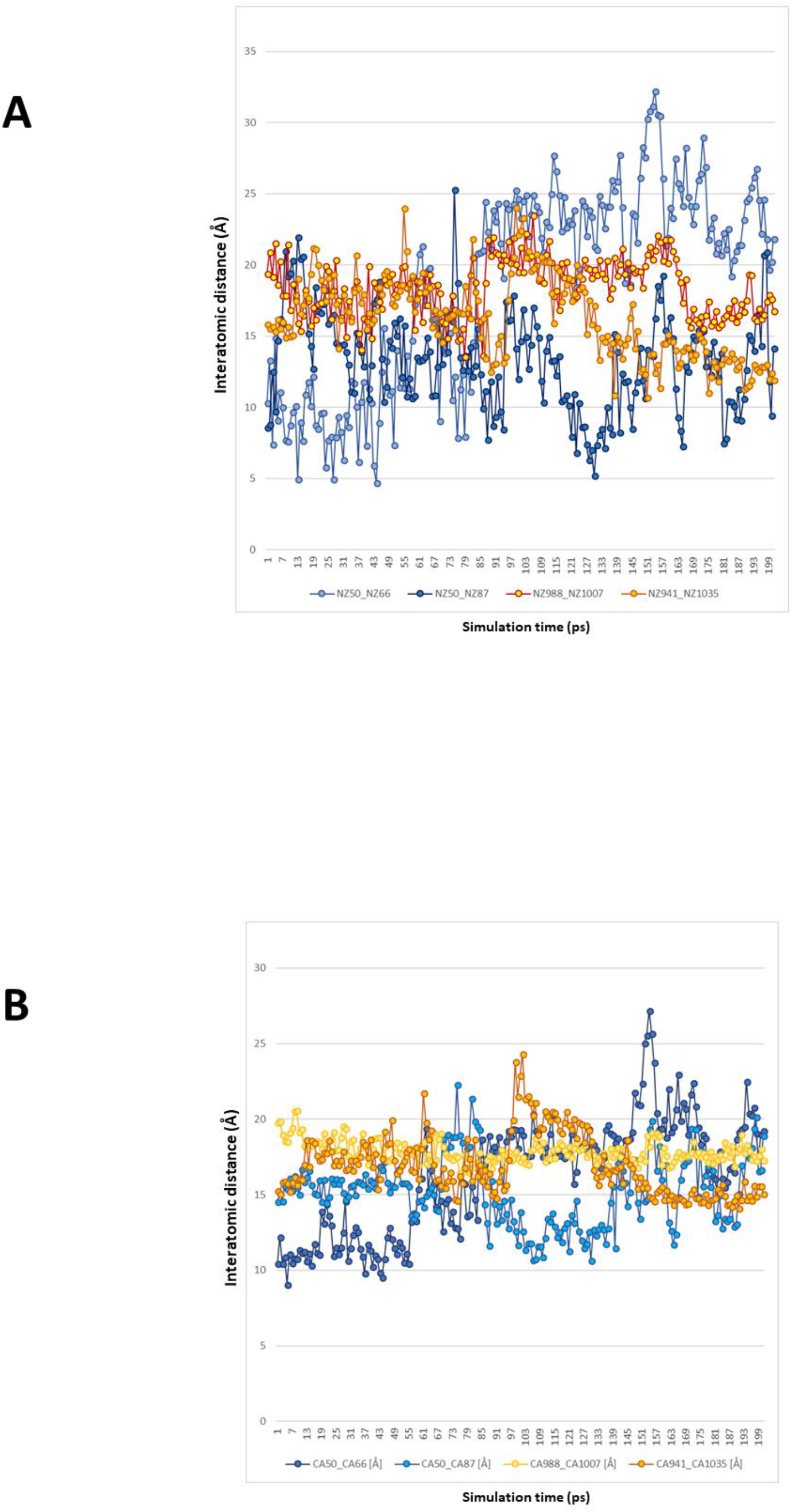
Comparison of the conformational dynamics of the NZ vs CA atoms in protein domains. **A.** NZ-NZ interatomic distances in the flexible domain (NZ50-NZ66, NZ50-NZ66) exhibit greater amplitude of molecular motion, compared to those in the rigid domain (NZ941-NZ1035, NZ988-NZ1007). **B.** CA-CA interatomic distances in the flexible domain (CA50-CA66, CA50-CA87) exhibit greater amplitude of molecular motion, compared to those in the rigid domain (CA941-CA1035, CA988-CA1007).

## Discussion

Our extensive experimental and computational analysis of XL-MS methodology applied to structural interrogation of a multi-subunit bacterial RNAP core provided a number of important insights which can lead to further improvement and broader application of this approach. We have reported a 44.9 % increase in the depth of XL-MS analysis of the already highly efficient NHS-ester-based X-linking by introducing 2 additional reagents (DSM4 and DSSeb(DSM8) in additional to the widely used DSS (DSM6). The addition (DSM8) or removal (DSM4) of 2 methylene groups from the X-linker spacer (relative to DSS), responsible for this increase, did not require modifications to the experimental conditions or sample processing (except for the straightforward alterations of X-link search parameters). It bears noting that DSM4 and DSM8 are now either commercially available, or can be economically made to order at a proteomics grade, allowing for immediate and broad deployment of this approach.

This depth of X-linking was achieved without any enrichment for X-linked peptides, apart from the data-dependent selection of ions charged at ≥4 during LC-MS/MS. Incorporation of the orthogonal methods of enrichment, such as the use of multiple proteases or the off-line fractionation by strong cation exchange and size exclusion peptide chromatography, would lead to further improvements of depth of coverage and X-link discovery rates^48-50^, especially for high complexity or low abundance *in vivo* samples.

The quantitative exploration of DSM4, DSM6, and DSM8 conformational mobility in MD simulations revealed the basis of the substantial overlap in the sets of non-redundant X-links formed by each reagent, and the generation of X-links unique to each set. Atomic trajectories in each simulation also allowed us to offer quantitative estimates of the X-linker-specific distance restraints (their median lengths, as well as the upper and lower limits) to replace currently underperforming estimates, based solely on the Euclidean length of the X-linker. Using extensive datasets (~5000 precursors each) of high-confidence X-links we have discovered that the shorter X-linker generated data skewed towards a few extremely abundant X-links, thereby reducing the overall number of non-redundant entries in each individual experiment. This finding is particularly important for XL-MS interrogation of high-value/low-abundance samples. Whenever the sample scarcity precludes an extensive optimization of the experimental conditions, longer X-links are more likely to produce a greater number of non-redundant X-links than the shorter ones (DSM8>DSM6>DSM4) (VS and EN, unpublished observations).

Explicit MD simulations of thermal motions of the experimentally X-linked residues provided realistic, quantitative estimates of the fold-dependent distance restraints. We were able to demonstrate that residues embedded into the flexible, unstructured loop regions undergo motions, vastly exceeding those of residues from the more compact domains. Together these findings indicate that, in order to improve the quality of X-links-guided protein-protein docking, the distance restraints involving residues from the flexible/highly mobile domains should be given less weight/lower violation penalties compared to the ones connecting residues from rigid domains. They also argue that use of CA-CA restraints substantially reduces the uncertainty in the data (thereby improving docking efficiency) compared to more relaxed NZ-NZ restraints (see **Supplementary Discussion** for more details).

## Acknowledgements

We would like to thank Zixuan Li for technical assistance and Tom Waltz for critical reading of the manuscript and invaluable insights. This work was supported by the NIH grant R01 GM107329 and by the Howard Hughes Medical Institute.

## Competing interests

None declared.

## Contributions

EN and VS conceived the study. VS carried out the experiments and data analysis under EN supervision. EN and VS wrote the manuscript.

## Materials and Correspondence

All correspondence and requests for materials should be submitted to Evgeny Nudler

## Materials and Methods

### Buffer components and consumables

Buffer components for cross-linking and protein purification were BioUltra grade (Millipore Sigma). LC-MS/MS was carried out with Thermo Scientific LC-MS grade reagents and solvents. Growth media components were from Thermo Fisher, protease inhibitor cocktail (ProBlock Gold Bacterial 2D) was from Gold Biotechnology. Low-binding pipet tips (Corning DeckWorks) and tubes (Protein LoBind, Eppendorf) were used throughout the experimental workflow.

### RNA polymerase expression and purification

RNA polymerase core was expressed in XJb(DE3) *E. coli* cells (ZymoResearch) and purified as previously described^1^, with additional polishing step: size exclusion chromatography on Superose Increase 6 10/300 GL (GE) in X-linking buffer (50 mM HEPES (pH 7.5), 500 mM NaCl, 2 mM MgSO_4_, 1 mM DTT). Fractions containing pure RNAP core (analyzed by SDS-PAGE) and exhibiting polydispersity less than 10% and the molecular weight estimate of 385-390 kDa (389 kDa theoretical) in dynamic light scattering experiments^2^ (DynaPro Nanostar (Wyatt Technologies)) were pooled together and used in XL-MS experiments.

### XL-MS: X-linking, sample processing and analysis

DSM4, DSM6, and DSM8 X-linkers (Proteochem) were dissolved in oxygen-depleted anhydrous DMSO (ZerO2, Millipore Sigma) at 100 mM stock concentration and added to 100 μl of RNAP core solution (0.2 mg/ml) in X-linking buffer to final concentration of 250-500 μM. X-linking was carried out for 45 min at 25°C in disposable cuvette (UVette, Eppendorf) with continuous monitoring of the polydispersity by dynamic light scattering to avoid aggregation of the sample (auto-attenuation of laser power, 10 acquisitions (5 sec each) per measurement). The reactions were quenched by addition of Tris-HCl (pH 7.5) to a final concentration of 10 mM.

Samples were dialyzed against 100 mM ammonium bicarbonate, reduced with 50 mM TCEP at 60°C for 10 min and alkylated with 50 mM iodoacetamide in the dark for 60 min at 25°C. Digestion was carried out at 37°C overnight with 0.5 μg sequencing grade modified trypsin (Promega) in 100 mM ammonium bicarbonate. The resulting peptides were passed though C18 Spin Tips (Thermo Scientific) before elution with 40 μL of 80% acetonitrile (ACN) in 0.1% trifluoroacetic acid. Eluted peptides were dehydrated in vacuum and resuspended in 20 μL 0.1% formic acid for MS analysis.

Peptides were analyzed in the Orbitrap Fusion Lumos mass spectrometer(Thermo Scientific) coupled to an EASY-nLC (Thermo Scientific) liquid chromatography system, with a 2 μm, 500 mm EASY-Spray column. The peptides were eluted over a 120-min linear gradient from 96% buffer A (water) to 40% buffer B (ACN), then continued to 98% buffer B over 20 min with a flow rate of 250 nL/min. Each full MS scan (R = 60,000) was followed by 20 data-dependent MS2 (R = 15,000) with high-energy collisional dissociation (HCD) and an isolation window of 2.0 m/z. Normalized collision energy was set to 35. Precursors of charge state 4-6 were collected for MS2 scans; monoisotopic precursor selection was enabled and a dynamic exclusion window was set to 30.0 s.

LC-MS/MS *raw* spectra were converted into *mgf* format using Proteome Discoverer (Thermo Fisher) and searched with pLink^3^. pLink *xlink.ini* file, containing X-linker search parameters, was amended to include DSM4=[K [K 110.0367 110.0367 128.0477 128.0477, DSM6=[K [K 138.068 138.068 156.079 156.079, and DSM8=[K [K 166.0993 166.0993 184.1103 184.1103. X-linked peptide search space was defined by combining *E. coli* RpoA (P0A7Z4), RpoB (P0A8V2), RpoC (P0A8T7), and RpoZ (P0A800) sequences into *RNAP.fasta* file. Search parameters were defined in the *pLink.ini* file by setting the enzyme name to trypsin, maximal number of missed cleavages to 3, maximal e-value to 0.001. Amino acid modifications were limited to 1 constant (Carbamidomethyl[C]), and 3 variable (Oxidation_M, Gln->pyro-Glu, and N-acetyl_Protein) ones. Samples of the *ini* and *fasta* files are included in the Supplement.

### Molecular dynamics simulations

In order to estimate the dynamic range of DSM4/6/8 spacers in solution we performed molecular dynamic simulation of each reagent using YASARA Dynamics^4^ (YASARA Biosciences GmbH). Starting structures in *pdb* format were generated from DSM4/6/8 structural formula, MD simulation was carried out for 25 ns in explicit water with 0.9% NaCl, at 298K in AMBER14 forcefield.

Starting *pdb* coordinates for simulations of molecular motions of RpoB residues 936-1046 and RpoC residues 36-103 were extracted from 4lk1. Simulations were carried out for 20 ns in the same conditions as described above. For detailed information on the settings and outcome of the simulations see YASARA *run* and *analysis* macros included in the Supplement.

